# GenOrigin: A Comprehensive Protein-coding Gene Origination Database on the Evolutionary Timescale of Life

**DOI:** 10.1101/2020.10.17.342022

**Authors:** Yi-Bo Tong, Meng-Wei Shi, Sheng Hu Qian, Yu-Jie Chen, Zhi-Hui Luo, Yi-Xuan Tu, Chunyan Chen, Zhen-Xia Chen

**Affiliations:** Hubei Key Laboratory of Agricultural Bioinformatics, College of Biomedicine and Health, Huazhong Agricultural University, Wuhan, Hubei 430070, PR China; College of Life Science and Technology, Huazhong Agricultural University, Wuhan, Hubei 430070, PR China; Center for Ecological and Environmental Sciences, Northwestern Polytechnical University, Xi’an 710072, PR China

## Abstract

The origination of new genes contributes to the biological diversity of life. New genes may quickly build their own network in the genomes, exert important functions, and generate novel phenotypes. Dating gene age and inferring the origination mechanisms of new genes, like primate-specific gene, is the basis for the functional study of the genes. However, no comprehensive resource of gene age estimates across species is available. Here, we systematically dated the age of 9,102,113 protein-coding genes from 565 species in the Ensembl and Ensembl Genomes databases, including 82 bacteria, 57 protists, 134 fungi, 58 plants, 56 metazoa, and 178 vertebrates, using protein-family-based pipeline with Wagner parsimony algorithm. We also collected gene age estimate data from other studies and uniformed the gene age estimates to time ranges in million years for comparison across studies. All the data were cataloged into GenOrigin (http://genorigin.chenzxlab.cn/), a userfriendly new database of gene age estimates, where users can browse gene age estimates by species, age and gene ontology. In GenOrigin, the information such as gene age estimates, annotation, gene ontology, ortholog and paralog, as well as detailed gene presence/absence views for gene age inference based on the species tree with evolutionary timescale, was provided to researchers for exploring gene functions.

## INTRODUCTION

Understanding the origination of genes is crucial to uncover the molecular mechanisms underlying the functional and phenotypic diversity (1). Gene origination has thus attracted wide interests from geneticists and evolutionists since the early twentieth century. Recently-formed young genes, like the first reported three-million-year-old gene *Jingwei* in *Drosophila* (2), were discovered, providing precious models to explore the evolutionary mechanisms of gene origination. So far, eleven mechanisms, among which gene duplication is considered to be a major player, have been found to be involved in gene origination (1,3).

The time of gene origination, namely, gene age, is closely related to its function (4). Gene age is correlated with the order of gene expression in the development of *Drosophila,* nematode, and zebrafish (1,5). Gene age is also correlated with diseases in human. Disease-causing genes tend to be old genes (6). Mutations in old genes may disrupt basic cellular functions such as genome integrity and transcriptional translation, while mutation in young genes may affect cell signaling and other functions (7). The origination of cancer genes corresponds to the emergence of multicellularity (8). The importance of gene age to functional study makes a comprehensive resource of gene age in great demand.

However, a comprehensive resource of gene age covering multiple species is still lacking. Although several studies have estimated gene age in certain species, most species have not been covered (9–11). Even in the covered species, the comparison of gene age inferred from various datasets was constricted due to the differences in annotations and methods among the studies. So far, there has been only one database on gene age estimates called GenTree (11), which focuses on human genes while provides gene age data of several other species including fly, mouse, rat, opossum, and chicken only for bulk download. Furthermore, the gene age estimates in GenTree are presented as branches, which depends on species used to build the phylogenetic tree, making it difficult to intuitively quantify the gene age estimates within a study or compare the gene age estimates across studies.

Three major challenges in providing a wide range of gene age assessments are the heterogeneity in the quality of genomes and annotations across species, the requirement for large computational cost, and the disagreement in gene age estimates among studies using different dating methods and ortholog datasets. First, the assembly level of genomes varies from contig to complete genome, and the annotation of model organisms are usually far more comprehensive than non-model organisms, restricting the unbiased comparative analysis (12). Second, the identification of orthologs, or genes in different species that descended from the same gene in their last common ancestor, is a prerequisite for gene age inference. However, complexity and computational requirements of ortholog identification from multiple species are substantial (13). Only few reference resources, like Ensembl Compara (14,15), explicitly provide orthology information (16). Third, two main gene-dating methods, protein-family based pipeline (FBP) and synteny-based pipeline (SBP), are optimal for either old genes or young genes (11). Also, the dating results rely on the ortholog datasets and the phylogenetic tree (17), making it impractical for the integration of gene age estimates from different resources.

To tackle the challenges and enable the query of gene age for a wide range of assessments across species, we developed the online database GenOrigin. It covered 9,102,113 protein-coding genes of 565 species from Ensembl (www.Ensembl.org, release 98) (18) and Ensembl Genomes (http://www.ensemblgenomes.org/, release 45) (19), which provide high-quality reference genomes and annotation for a wide range of species. GenOrigin systematically infers gene age using FBP with Wagner parsimony algorithm, based on the same timetree derived from TimeTree database (http://www.timetree.org) (20) and the orthology information from Ensembl Compara (14,15). With GenOrigin, researchers can easily query the age estimates, in terms of both million years and phylogenetic branches, of any gene in the 565 species and view its evolutionary trajectory, or download batches of gene age estimates for systematically studying the evolution and function of genes.

## METHODS

### Construction of timetree

We took species from Ensembl (www.Ensembl.org, release 98) (18) and Ensembl Genomes (http://www.ensemblgenomes.org/, release 45) (19) for timetree construction (Figure 1). Ensembl (18) and Ensembl Genomes (19) are central repositories of the vertebrate domain and five invertebrate domains (bacteria, protists, fungi, plants, and metazoa) with the same platform, providing genomes, gene annotations, and domain- specific comparative genomics information. The gene annotations they provide are evidence-based, updated regularly, and relatively consistent across all species. They also inferred pairwise relationships between homologs, diverged by a speciation event (namely orthologs) or by a duplication event within one genome (namely paralogs) (21,22), based on gene and species tree reconciliation (14,15). Homologous relationships of genes from different domains can be inferred through the mediation of representative species in the six domains involved in the Pan-taxonomic Compara (19). Therefore, Ensembl and Ensembl Genomes are optimal for systematically tracing evolutionary trajectory of genes in a wide range of species.

**Figure 1.**
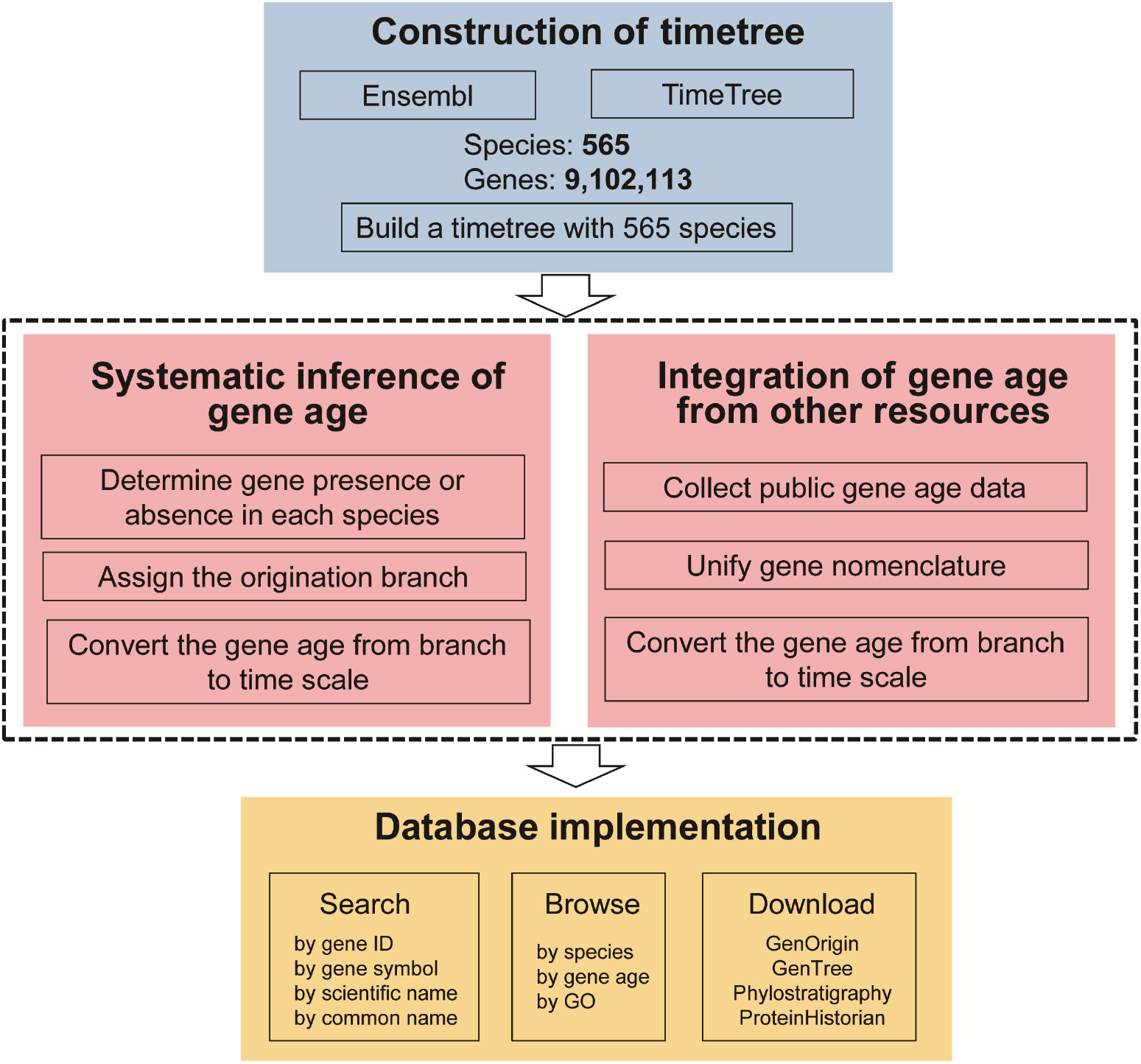
Workflow of GenOrigin construction. GenOrigin used 565 species from both Ensembl (www.Ensembl.org) and TimeTree (www.timetree.org) to build the timetree. It systematically inferred age of the 9,102,113 genes from the 565 species based on their presence/absence in each species on the timetree. Gene age data were also collected from other resources and integrated into GenOrigin. Users can search, browse, and download all the gene age data from GenOrigin.

We then used TimeTree database (20) to build the timetree, a phylogenetic tree with time scale (Figure 1). TimeTree assembles a tree of life with time scale in million years (myr) based on thousands of published studies, and can build a timetree with custom species list (20). Among the species from Ensembl and Ensembl Genomes, 565 species were included in TimeTree, and were used to construct the timetree. We downloaded the timetree file in newick format from TimeTree, and processed the file using Biopython (version 1.72) (15) module *Phylo* for further analysis.

### Derivation of gene families from Ensembl and Ensembl Genomes

We extracted the orthology information, including orthologs and paralogs, of all the protein-coding genes from protein-family-based Ensembl Compara data (14) downloaded in bulk via the Ensembl and Ensembl Genomes FTP server in TSV format. Ensembl Compara is the platform used by both Ensembl and Ensembl Genomes to offer cross-species resources and analysis of gene families. Ensembl Compara inferred domain-specific or pan-taxonomic orthologs of all the protein-coding genes based on both similarity and the reconciliation of gene and species trees. The pipeline includes classifying the representative protein sequences into their families using an HMM search on the TreeFam HMM library (23), clustering the orphaned genes into additional families based on the Blast e-values (24), aligning each family using M-Coffee (25) or Mafft (26), building gene trees for the families, and conciliating the gene trees against the species trees to infer gene pairwise relations of homology types (15). It can find one-to-one, one-to-many and many-to-many orthologs between species within each domain or involved in the Pan-taxonomic Compara (14).

For each gene of any species in a domain, we retrieved its orthologs in the domain’s Compara, and expanded its distribution to Pan-taxonomic Compara by the mediation of the closest neighbour species in Pan-taxonomic Compara with its ortholog. We further expanded its distribution to other five domains by the mediation of the species, with its most similar ortholog, shared by Pan-taxonomic Compara and Compara of the five domains. All the orthologs of each gene were kept for the inference of gene age.

### Systematically dating protein-coding genes on the branches of the timetree

We used *Count* (27) to infer the branches where gain and loss of genes occurred in the timetree based on the presence and absence of genes (orthologs) in each species (Figure 1). *Count* implemented Wagner parsimony and Dollo parsimony, two widely used ancestral history reconstruction algorithms. In contrast to Dollo parsimony, which assumes a single gain event of each gene family, Wagner parsimony allows multiple gain and loss events (28). To prevent false positives in gene presence due to horizontal gene transfer from inflating age distributions, we computed Wagner parsimony using AsymmetricWagner application of *Count*. Its option for relative penalty of a gain with respect to loss (-gain) was tuned from 0 to 2.5 with a step of 0.1, and set to 1.4 when the consistency with GenTree was over 60% for the total inferred human gene age and reached maximum for genes younger than 435 myr (corresponds to the divergence of vertebrate ancestors, and the earliest divergence time of species used in GenTree).

The output of *Count* were analysed to infer the age of each protein-coding gene with a modified pipeline (9–11) using customized Python scripts. In brief, we assigned levels to the timetree branches based on their hierarchic rank from the root node to the terminal nodes, and referred to the branch started at the root node as branch 0 (the lowest level). The evolutionary trajectory of each gene was traced from the terminal node (species) to the root node until a loss event was encountered in branch N, and the gene would be dated to branch N-1. When there was no loss event encountered, the gene would be dated to branch 0.

### Inference of young gene origination mechanisms

We modified a previous pipeline (9–11) to infer the potential origination mechanisms, including de novo origination, retroposition, retroposition like, DNA-based duplication, or DNA-based duplication like, for young genes <100 myr, whose clues based on paralog comparisons were relatively clear. Young genes without paralogs were assigned to de novo origination. The paralogs would be assumed to be the potential parental genes. Young genes that had only one coding exon while their paralogs had multiple coding exons were assigned to retroposition (identity >= 0.5) or retroposition like (identity is < 0.5). Young genes that had multiple coding exons and had paralogs with multiple coding exons were assigned to DNA-based duplication (identity >= 0.5) or DNA-based duplication like (identity is < 0.5). When young genes and their paralogs all had one coding exon, the young genes were assigned to DNA-based duplication (identity >= 0.5) or DNA-based duplication like (identity is < 0.5).

### Integration of gene age estimates from other resources

We also collected the gene age data from other resources and integrated them into GenOrigin with unified gene nomenclature (Ensemble ID) (Figure 1). UniProt IDs were converted into gene symbols through the API provided by UniProt (https://www.uniprot.org), and gene symbols were converted to Ensembl ID through Ensembl BioMarts.

We not only compared gene age estimates in branch-scale, but also normalized the gene age estimates from different resources to our timetree in time-scale. To convert the origination time from branches into molecular time estimates, the timetree was rebuilt for each study on different species, and used for dating gene age. The mid-point of the origination branch in myr was defined as the gene age.

### Gene expression analysis

We downloaded RNA-seq datasets from EBI ArrayExpress (www.ebi.ac.uk/arrayexpress) for gene expression analysis under accession number E-MTAB-6814, E-MTAB-6798 and E-MTAB-6769. The datasets covered the developmental stages from organogenesis to adulthood of seven major organs (i.e. brain, cerebellum, heart, kidney, liver, ovary and testis) in human, mouse and chicken from the same study (29). We used snakePipes (v1.3.0) (30), a workflow package for processing high throughput data, to estimate the gene expression level across tissues in the three species. In RNA-seq analysis, SnakePipes integrated STAR (v2.6.1) for read mapping, and featureCounts (v2.0.0) (31) for read quantification. We measured gene expression in TPM (Transcripts Per Kilobase Million) (32) based on read counts. The median TPM of all the developmental stages of a tissue in a species was taken as the expression level of the tissue in the species. Tissue specificity index tau was applied to measure tissue specificity of each gene (33).

### Database implementation

The GenOrigin database was constructed under the Flask open source framework (http://flask.pocoo.org/). All data were integrated into MongoDB (version 3.2.11). The web interface was designed and implemented using AngularJS (version 1.6.9) and was improved with some AngularJS libraries and several JavaScript libraries for more convenient use. The web browser Google Chrome is recommended for accessing GenOrigin.

## RESULTS

### The coverage and quality of gene age estimates in GenOrigin

As a comprehensive gene age database across species, GenOrigin provided users with its own gene age estimates in species from Ensembl (18) and Ensembl Genomes (19) using FBP (Figure 1), along with those from three widely used gene age datasets GenTree (11), Phylostratigraphy (8) and ProteinHistorian (28). We built a timetree using TimeTree (20) with 565 species, including 178 vertebrates from Ensembl, and 82 bacteria, 57 protists, 134 fungi, 58 plants and 56 metazoa from Ensembl Genomes, whose divergence time ranged from 0 to 4,290 myr. Gene ages were inferred for the 9,102,113 genes from 565 species, far more than the other three resources (1~33 species), in Ensembl and Ensembl Genomes (Table 1). The genes with inferred age in GenOrigin included 1,897,157 genes (20.8%) younger than 100 myr, and 1,165,372 genes (12.8%) older than 1,000 myr. Specifically, in humans, 5,485 genes (23.7%) and 3,554 genes (15.4%) were younger than 100 myr and older than 1,000 myr.

**Table 1.**
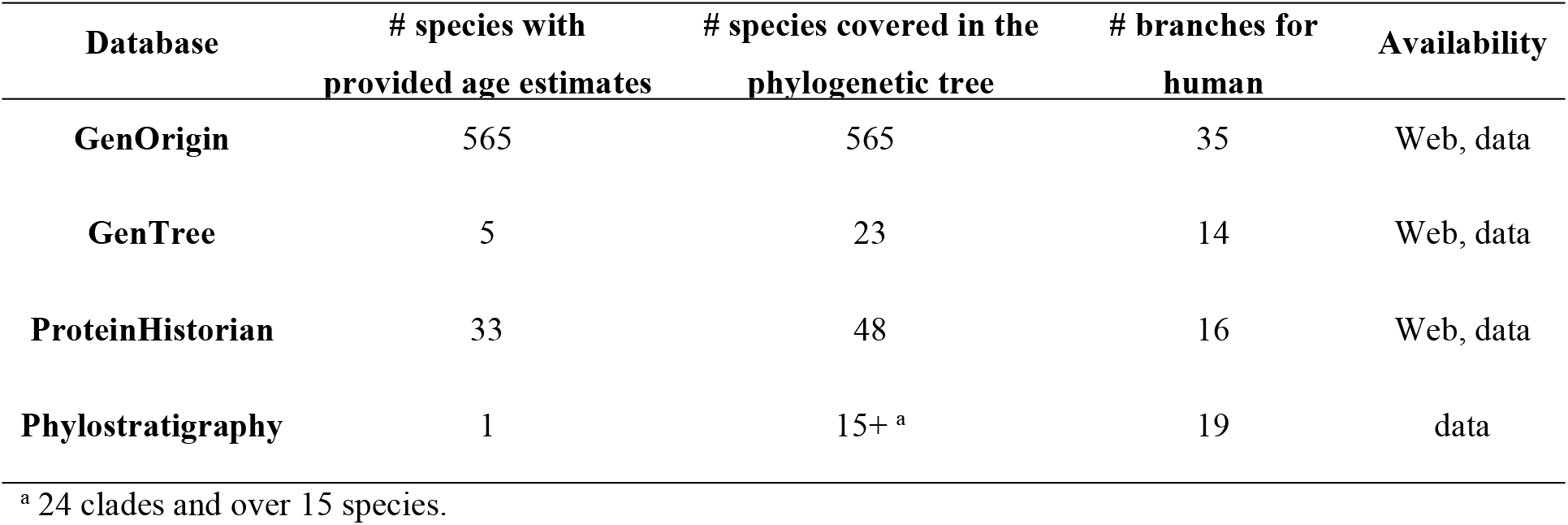
Species coverage of the major resources providing orthology

To validate the quality of our inferred gene ages, we compared human gene ages in GenOrigin with GenTree (11), Phylostratigraphy (8) and ProteinHistorian (28). GenTree is a recently published resource for human gene age using the conservative SBP optimal for young genes (11), while Phylostratigraphy and ProteinHistorian are another two datasets using the FBP with Dollo parsimony (8) and Wagner parsimony (28), respectively. The evolutionary trajectory from root to human included 35 branches for GenOrigin, more than GenTree (14 branches), Phylostratigraphy (19 branches) and ProteinHistorian (16 branches), increasing the accuracy of human gene age estimates. For example, the gene cyclin dependent kinase 9 (CDK9, ENSG00000136807) was dated to 1480-1496 myr by GenOrigin, while >435 myr by GenTree, >4290 myr by Phylostratigraphy, and 1105-1496 myr by ProteinHistorian.

We measured the consistency of age estimates for genes commonly covered in GenOrigin and other three datasets, and found GenOrigin was consistent with GenTree for 11,753 genes (64.2%), while consistent with Phylostratigraphy and ProteinHistorian for less than 30% genes (Supplementary Figure S1). The differences between GenOrigin and other datasets might be resulted from the differences in phylogenetic tree, gene annotation, orthology information, and dating method. Among the genes consistent with GenTree, 11,126 (94.7%) were at the branch 0 of GenTree (>435 myr), which corresponded to branch 0-16 of GenOrigin, branch 0-12 of Phylostratigraphy and branch 0-6 of ProteinHistorian, demonstrating that the lower consistency with Phylostratigraphy and ProteinHistorian than GenTree was mainly resulted from their higher resolution of gene estimates for genes >435 myr.

To separate the effects from branch differences among datasets, we dated human gene ages based on the same species used in GenTree with GenOrigin pipeline. We found GenOrigin was highly consistent with GenTree, Phylostratigraphy and ProteinHistorian for 61.2% to 81.6% of all the genes (Figure 2A), similar as the consistency of GenTree with Phylostratigraphy and ProteinHistorian (~60%-70%) (11), demonstrating at least comparable, if not better, performances of GenOrigin in gene age estimation with other FBP resources.

**Figure 2.**
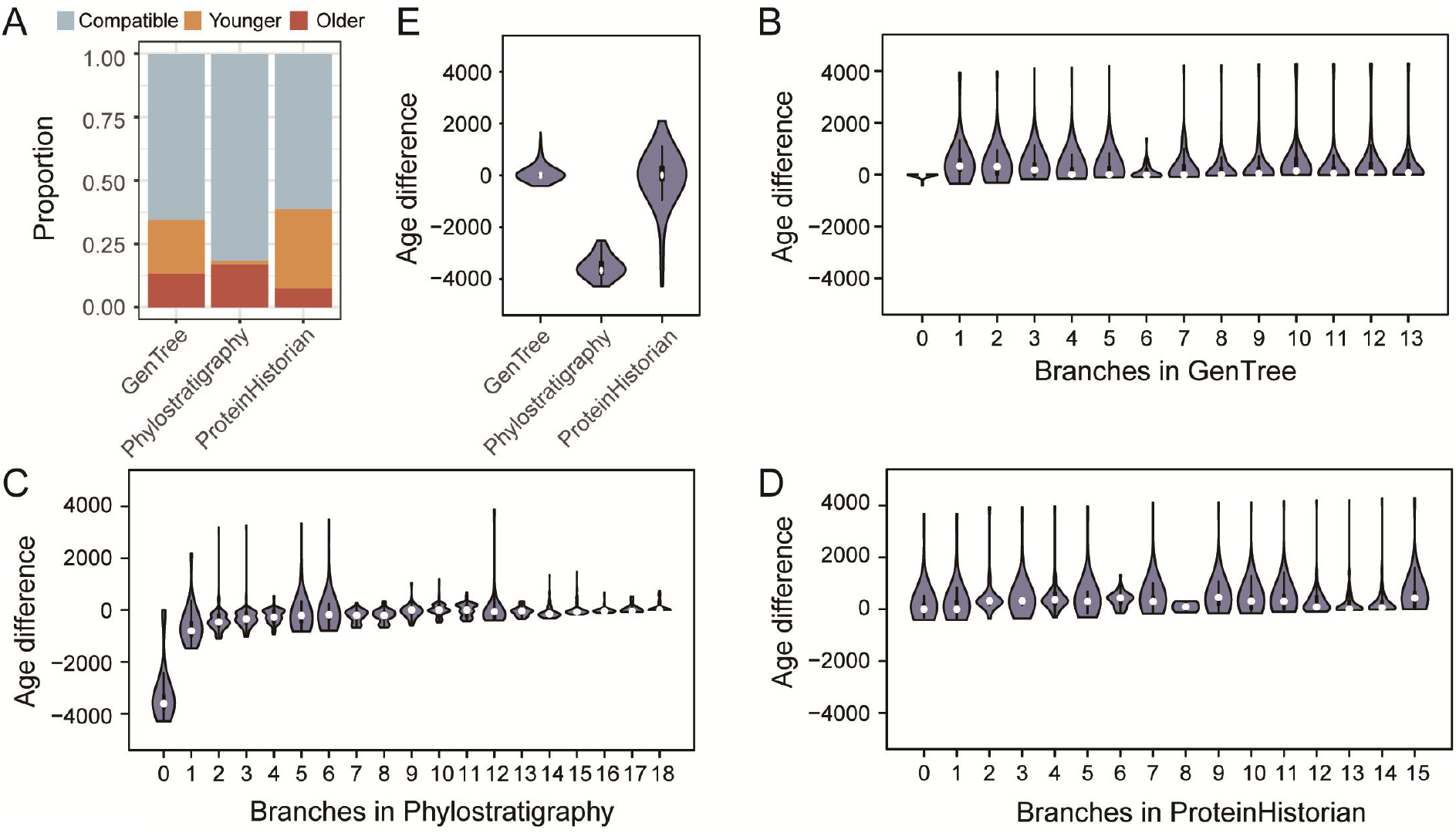
Comparison of gene age estimates in GenOrigin with GenTree, Phylostratigraphy and ProteinHistorian. (A) Comparison of age estimates in GenOrigin with those in other datasets based on the phylogenetic tree built with their shared species. Compatible: equal to GenOrigin; Older: Older than GenOrigin; Younger: Younger than GenOrigin. (B-D) Age difference (myr) between GenOrigin and GenTree (B), Phylostratigraphy (C) and ProteinHistorian (D). GenOrigin gene age estimates are older than other datasets when age difference >0, and younger when age difference <0. (E) Age difference between GenOrigin and other datasets for genes in the branch 0 and branch 1 of Phylostratigraphy.

We further measured the inferred age difference between GenOrigin and other datasets, and found they were all centralized around 0 except for the oldest genes in the branch 0 and branch 1 of Phylostratigraphy (Figure 2BCD). We then compared the inferred age difference of genes in the branch 0 and branch 1 of Phylostratigraphy between GenOrigin and GenTree/ProteinHistorian, and found they were both centralized around 0 (Figure 2E), supporting the overestimation of gene age for Phylostratigraphy (11,28,34) instead of the underestimation of gene age for GenOrigin.

### Applications of GenOrigin

The gene age estimates of 565 species provided by GenOrigin based on the same timetree enables users to query the approximate origination time of genes, and use it as a reference for a wide range of assessments.

As an example, we investigated the distribution of gene age in human, mouse and chicken, and found a common peak in all the species at 645 myr (Figure 4A), which corresponds to the occurrence of two whole genome duplications before the vertebrates radiation (615-684 myr) (38), supporting that duplication is the predominant mechanism of new gene origination.

We then assessed the evolution of basic genomic features in the three species over time. As to gene length, we found that the younger genes were shorter than older genes in all the species (Spearman’s rho: 0.80~0.86, p<0.001) (Figure 4BCD), which was consistent with previous study in human, stickleback and zebrafish (39). We further looked into average exon length and exon number, and found that they both positively correlated with gene age estimates (Spearman’s rho: 0.69~0.83 for exon length and 0.89~0.93 for exon number, p<0.001) (Figure 4EF), contributing to the positive correlation between gene age and gene length.

We also studied the relation of gene age estimates with expression in the three species based on the RNA-seq data across developmental stages and tissues of the three species from the same study (29). We found that younger genes had the lower expression level (Spearman’s rho: 0.92~0.95, p<0.001) (Figure 4E) and higher tissue specificity (Spearman’s rho: −0.82~ −0.76, p<0.001) (Figure 4F) in all the three species. Considering expression quantification of duplicated genes may be problematic due to multi-mapped reads, we further remained only genes without paralogs for the same analysis, and found similar pattern (Supplementary Figure S2, S3). This pattern is consistent with previous study based on SBP in human and mouse (40), suggesting a general pattern that new genes might need time to integrate into the pre-existing network and acquire more interactions (41).

These similar evolutionary patterns of genomic features as previous study using different dating pipeline in the same and different species validated the patterns in a broader range, and thus demonstrating our strength in comparative study across multiple species. The generality of the pattern provides firmer foundation for further exploration of underlying biology.

### Browse, search and download of the database

GenOrigin is a user-friendly gene age database from which users can search, browse, and download gene age data of 565 species (Figure 3A). Search can be conducted by using keywords, such as Ensembl gene ID, gene symbol, species scientific name, species common name, and GO term/name (Figure 3B). Users can search for one or a class of genes with a complete (e.g. “SESN1” as gene symbol for exact match) or partial entry (e.g. “SESN” as gene symbol for partial match) at a time. The default is searching with “Exact Match”. Users can also search with partial entry by selecting “Partial Match”.

**Figure 3.**
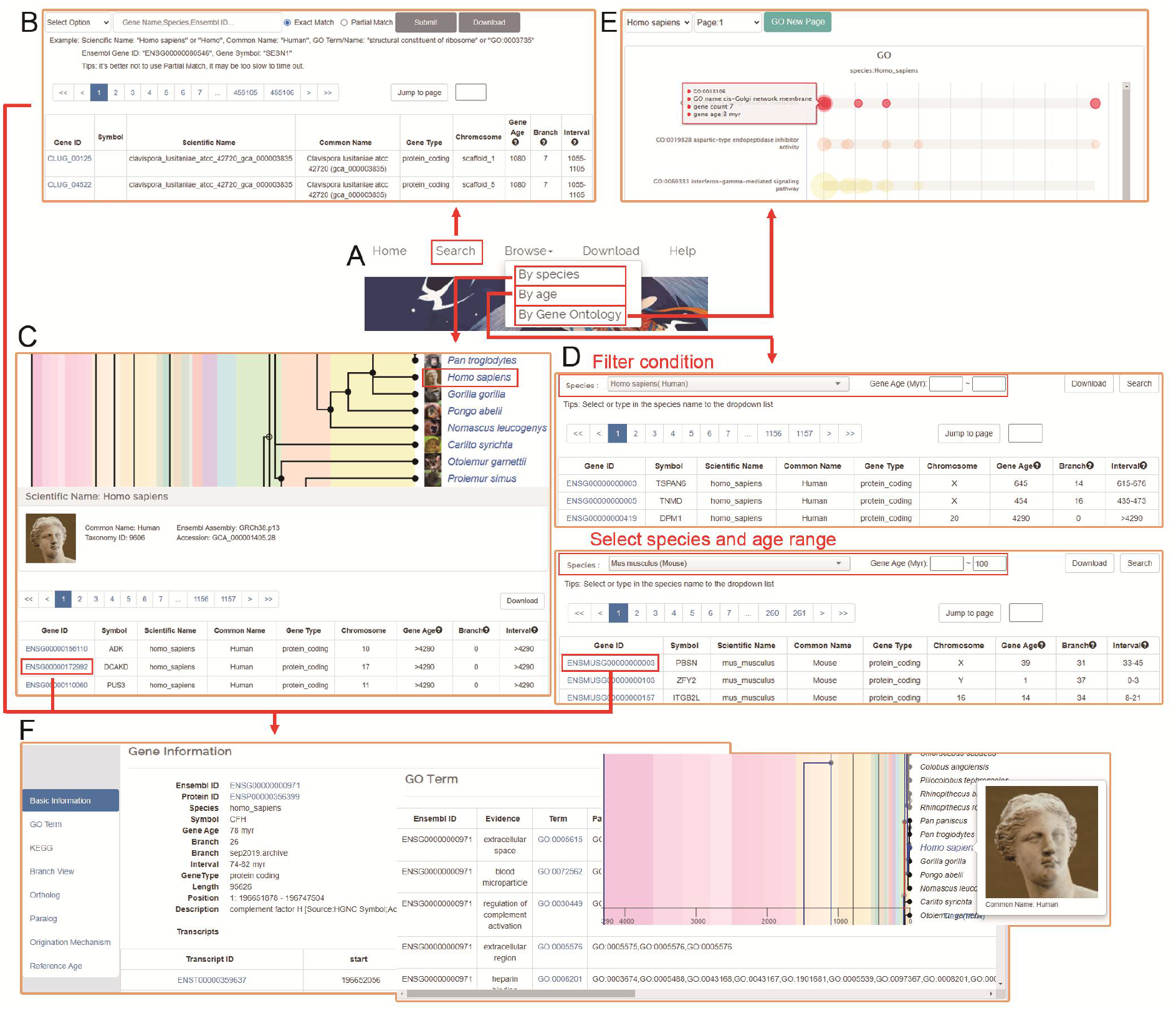
An overview of GenOrigin. (A) The homepage of GenOrigin. (B) Search by Ensembl gene ID, gene symbol, species scientific name and species common name. (C) Browse by species. The species tree was plotted by TimeTree (www.timetree.org) (D) Browse by gene age. (E) Browse by Gene Ontology. (F) Information of each gene, including basic information, GO, KEGG, branch view, ortholog, parolog and origination mechanism and reference age estimates. The continuous blue line in the branch view represents the evolutionary trajectory from root to a particular species; the red branch represents the origination branch of the gene; the presence/absence of a gene in each species and branch is marked in black/gray nodes and branches, respectively.

**Figure 4.**
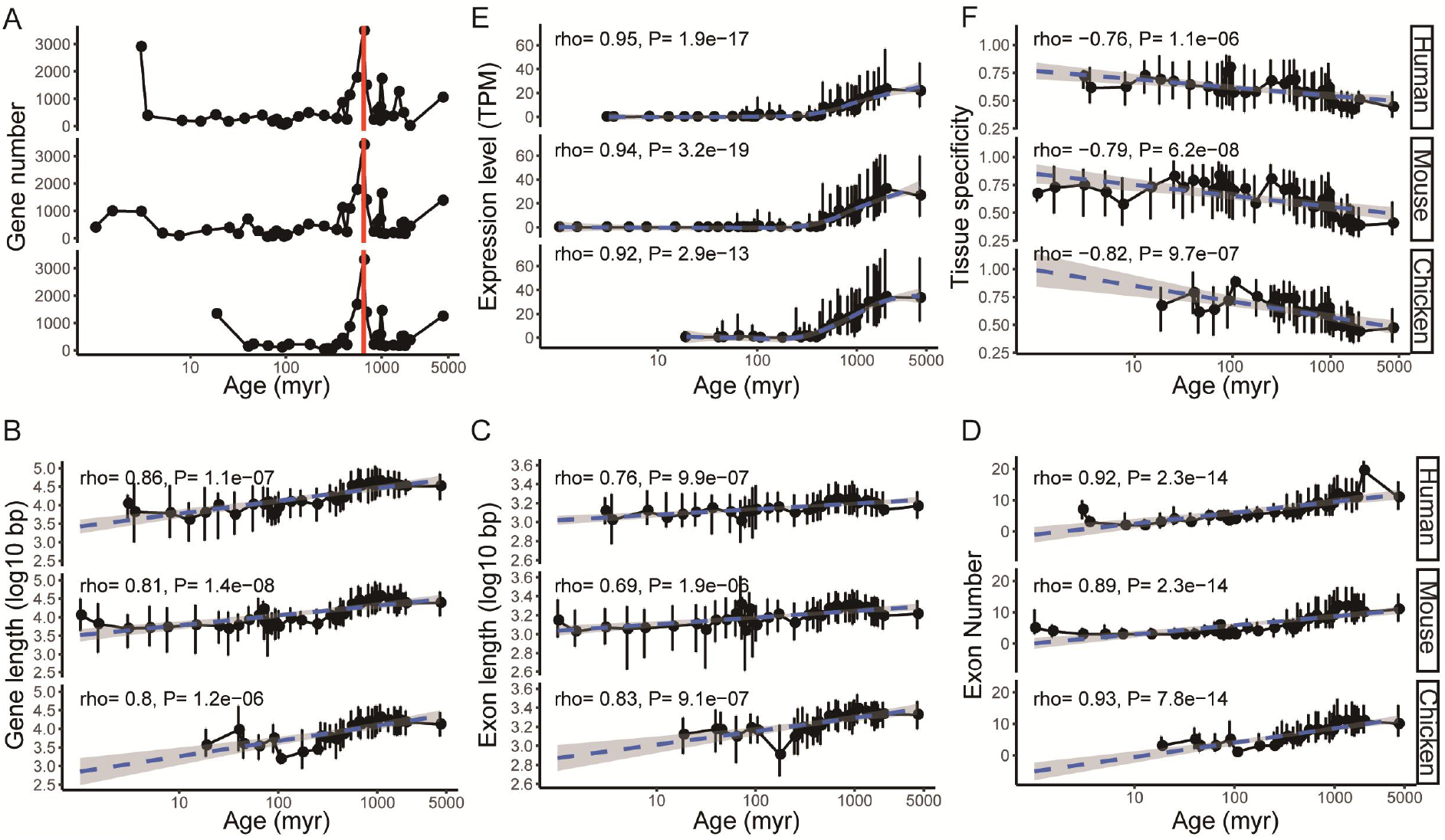
Application examples for using GenOrigin as a reference of gene age estimates for a wide range of assessments in human, mouse and chicken. (A) The distribution of gene age estimates based on the midpoint of inferred origination branch. The red line indicates the peak of gene number at 645 myr. (B-D) The relationship between gene age estimates and genomic features, including gene length (B), exon length (C), and exon number (D). (EF) The relationship between gene age estimates and expression patterns, including expression level (E) and tissue specificity (F). The blue dashed lines and shaded regions represented the fitting curves and their 95% confidence interval. The p value and correlation coefficient for Spearman’s correlation were shown on the top left.

Three browse modes, by species (Figure 3C), by gene age (Figure 3D), and by Gene Ontology (GO) (35) (Figure 3E), are provided. When browsing by species, users can pick a species of interest by selecting in the drop-down menu, or by clicking at the species name on the timetree (Figure 3C). When browsing by gene, users can select the species from the drop-down menu, and refine the results with the range of gene age (Figure 3D). For example, if users want to browse the young human genes originated in the recent 50 myr, they only need to select “Homo sapiens (Human)” in the drop-down menus of “Species” on the top left, enter “0” and “50” in the text boxes of “Gene Age (Myr)” on the top middle, and click the “Search” button on the top right. The search results will be presented in a table that contains gene age estimates in branch (on the “Branch” column) and in time-scale (mid-point of the origination branch on the “Gene Age” column, and time range of the origination branch on the “Interval” column) (Figure 3D). When browsing by GO, users can select the species from the drop-down menu to view the age distribution of GO-related genes, and retrieve genes related to a certain GO term by clicking at the bubbles. (Figure 3E).

All the search and browse results are directed to gene information pages (Figure 3F). We integrated gene annotation information and gene origin information inferred or collected by GenOrigin. Gene annotation information included the basic information, homologs and GO from Ensembl FTP, and KEGG (36) from Ensembl BioMart (37). The gene origin information inferred by GenOrigin included gene age estimates, branch view and origination mechanism. We displayed the gene age inference with an interactive timetree. On the timetree, the presence/absence of a gene in each species or branch was marked in black/gray, and the evolutionary trajectory from the root node to terminal node (the species of the gene) was highlighted in blue. Specially, the origination branch of the gene was labelled in red. Users could examine each branch by moving the mouse cursor to it (Figure 3F).

All the search results can be downloaded as CSV files for customized analysis by clicking the “Download” button on the top right. Alternatively, the gene age data of each species can be batch downloaded from the Download page.

## DISCUSSION

### GenOrigin is a comprehensive resource of gene age estimates

GenOrigin aims to provide users a comprehensive resource of gene age estimates by systematic gene age inference across 565 species and by integration of gene age inference from other resources. A comprehensive resource of gene age estimates requires wide coverage of genes, precise gene age estimates, unified dating pipelines, comparable age measurement among genes, as well as the integrated gene age data from multiple resources. GenOrigin attempts to achieve the goals: 1) It widely covers 9,102,113 genes from 565 species, more than GenTree (6 species), Phylostratigraphy (1 species) and ProteinHistorian (32 species), enabling users to retrieve gene age estimates of more species. 2) It involves 565 species diverged from each other 0 - 4290 myr ago on the timetree for the inference, which makes high resolution gene age estimates possible. As for human, the time range for the origination branch can be as small as 2.41 myr. 3) The same timetree is used to systematically date the gene age of the 565 species. The unified gene nomenclature and annotations (Ensembl, version 98; Ensembl Genomes, version 45) (18), as well as the standard dating pipelines, make it possible to compare gene age characteristics across species. 4) The origination time in branch is converted into myr based on the time scale from TimeTree (20). Such conversion makes it easy for users to compare the age across genes and datasets of a species. 5) We incorporate gene age data from three other datasets and ours so that users can check the consistency of the gene age estimates among datasets and choose gene age estimates from one of the datasets depending on their needs.

### Caveats

The inference of gene age is a complex problem relying on the validity and precision of the timetree, the completeness and accuracy of ortholog identification, and the effectiveness of the dating algorithms. Although we built the dating pipeline with the widely used resources and methods, we are expecting better and better performance of our dating results with the development of the resources and methods. For example, only annotated genes can be compared and thus missing genes may influence our age estimates. Although Ensembl (18) and Ensembl Genomes (19) have created a comprehensive and consistent baseline of reference annotation across species and plateaued on protein number for key species like human (Supplementary Figure S4) (42), their gene annotation is still requiring improvement, especially for non-model species. Also, the gene age inferred by GenOrigin is based on the orthology information of Ensembl Compara (14,15), which perform great for old genes but not for new genes. Meanwhile, for human genes, GenTree (11) integrates age inferred with functional genomic data using 18 vertebrate species diverged from each other 6.65 – 435 myr ago, thus, it is desirable for studying new genes, while not for investigating the genes older than 435 myr. Therefore, users should be cautious in dataset selection when querying the gene age estimates.

### Conclusion

In summary, GenOrigin is a comprehensive and user-friendly resource of gene age estimates covering the protein-coding genes from 565 species. It provides gene age systematically inferred by GenOrigin, along with gene age estimates integrated from three other resources, and it will contribute to the evolutionary and functional studies of genes and genomes.

## ACKNOWLEDGEMENT

We gratefully acknowledge Manyuan Long from the University of Chicago and Yong E. Zhang from Institute of Zoology, Chinese Academy of Sciences for the helpful suggestions. We thank Zhongliang Xue for providing the *jingwei* origin figure in the web page. We are also grateful to our users and all members in our lab for their valuable suggestions and comments.

## Supplementary Material

**Supplementary Figure S1.** Comparison of gene age estimates in GenOrigin with GenTree, Phylostratigraphy and ProteinHistorian based on the phylogenetic tree built with their own species.

**Supplementary Figure S2.** The relationship between gene age estimates and expression level in human, mouse and chicken for genes without paralogs. The blue dashed lines and shaded regions represented the fitting curves and their 95% confidence interval. The p value and correlation coefficient for Spearman’s correlation were shown on the top left.

**Supplementary Figure S3.** The relationship between gene age estimates and tissue specificity in human, mouse and chicken for genes without paralogs. The blue dashed lines and shaded regions represented the fitting curves and their 95% confidence interval. The p value and correlation coefficient for Spearman’s correlation were shown on the top left.

**Supplementary Figure S4.** The number of protein-coding genes in Ensembl version 92 to 100.

