## Supplementary figures and images for "GenOrigin: A Comprehensive Protein-coding Gene Origination Database on the Evolutionary Timescale of Life"

### Supplementary Figure S1

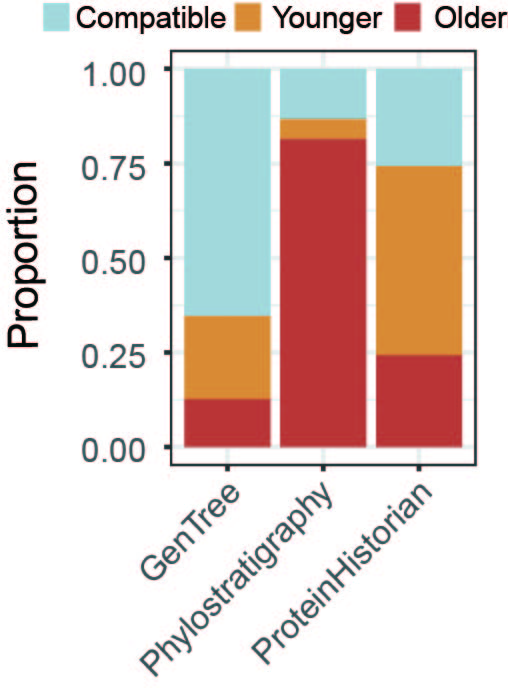

### Supplementary Figure S2

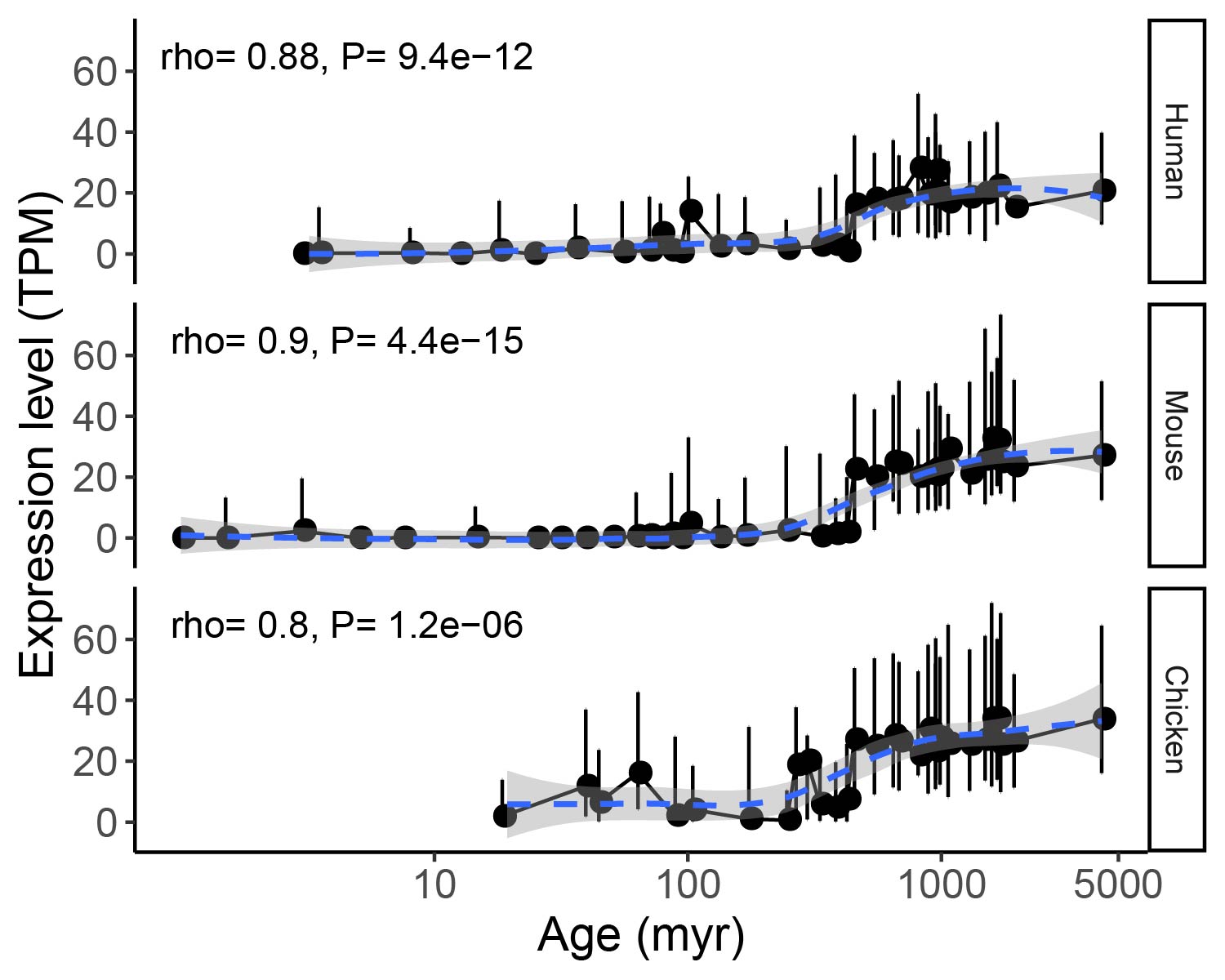

### Supplementary Figure S3

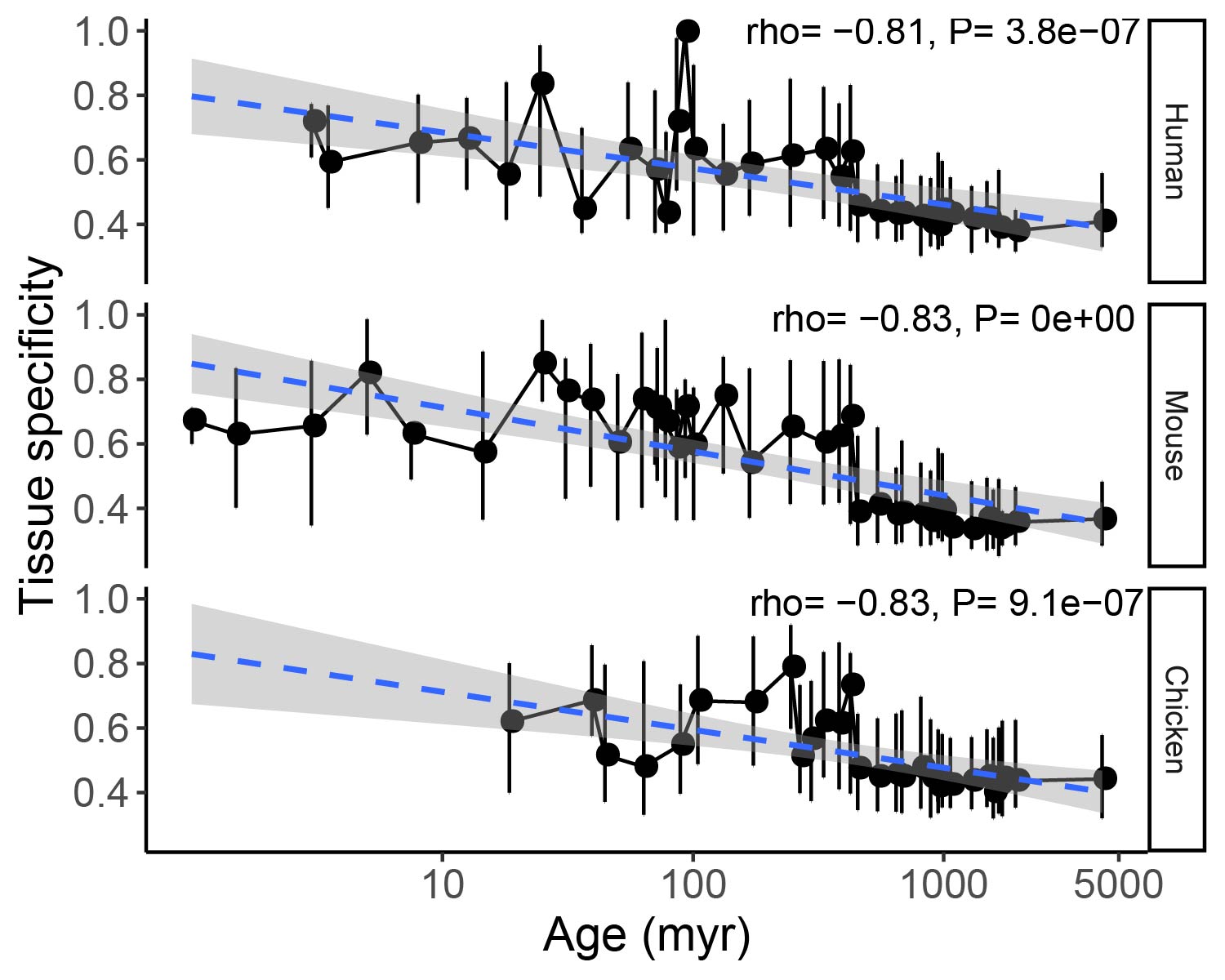

### Supplementary Figure S4

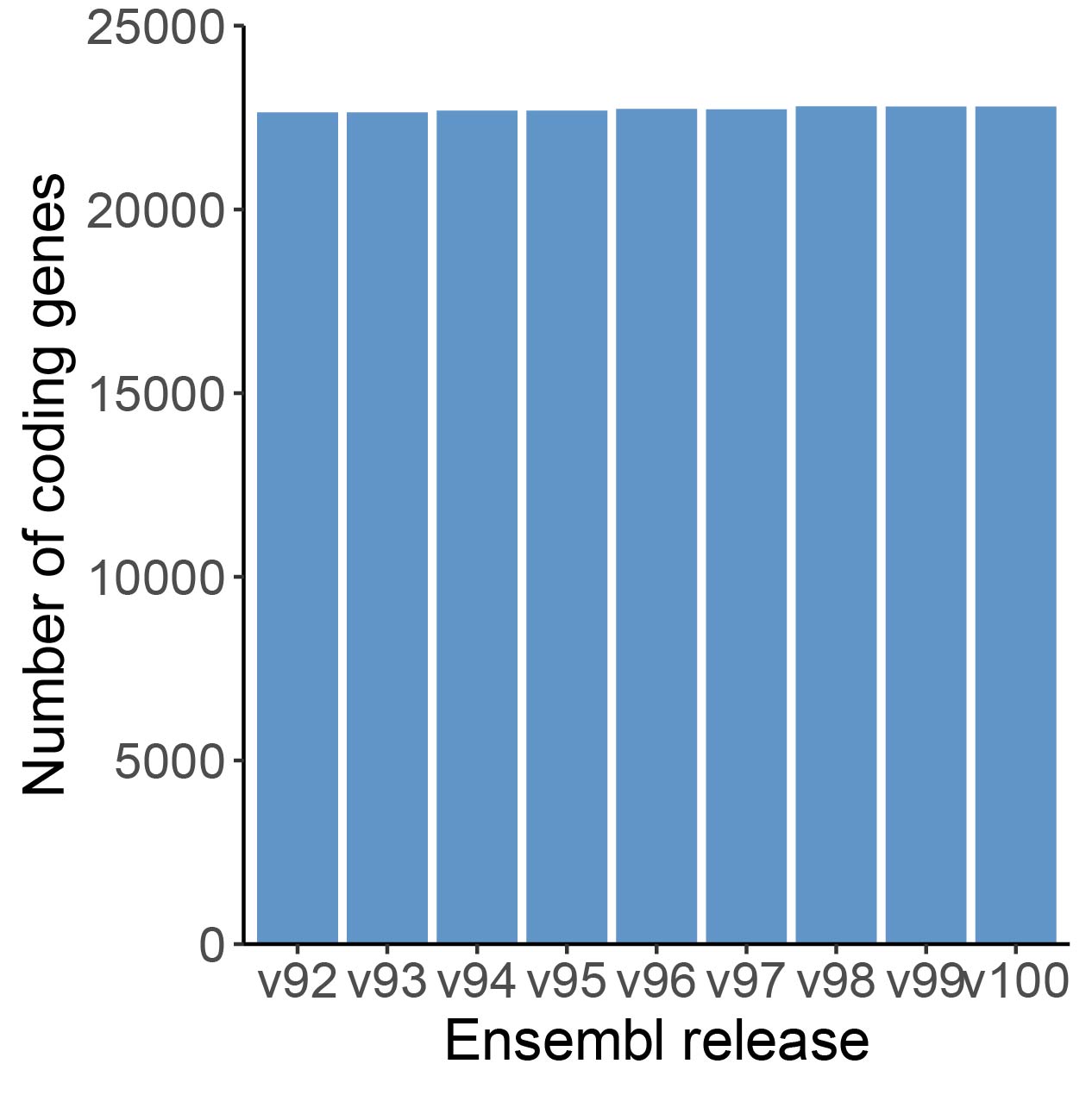
